# Unveiling how herpetofauna cope with land-use changes— insights from forest-cashew-rice landscapes in West Africa

**DOI:** 10.1101/2024.06.03.596777

**Authors:** Francisco dos Reis-Silva, Cristian Pizzigalli, Sambu Seck, Mar Cabeza, Ana Rainho, Ricardo Rocha, Ana Filipa Palmeirim

**Affiliations:** Global Change and Conservation Research Group, Faculty of Biological and Environmental Sciences, University of Helsinki, Viikinkaari 1 P.O. Box 65 00014, Finland; CIBIO/InBIO-UP, Research Centre in Biodiversity and Genetic Resources, University of Porto, Vairão, 4485-661, Portugal; Federação KAFO, Guiné-Bissau CP 1186, Centro Camponês de Djalicunda, sector de Mansaba, região de Oio, Guinea-Bissau; Centre for Ecology, Evolution and Environmental Changes & CHANGE - Global Change and Sustainability Institute. Departamento de Biologia Animal, Faculdade de Ciências, Universidade de Lisboa, 1749-016 Lisboa, Portugal; Department of Biology, University of Oxford, 11a Mansfield Rd, OX1 3SZ, Oxford, UK

**Keywords:** amphibians, agriculture, agroecosystems, Guinea-Bissau, habitat conversion, species diversity, reptiles, tropical forest

## Abstract

Agricultural-induced land-use change comprises a key driver of biodiversity loss across tropical forests. Guinea-Bissau, in West Africa, was formerly occupied by native forest-savanna mosaics. While savannas have long gave place to traditional rice agroecosystems, forests are now being transformed into cashew monocultures at unprecedented rates. The ecological impact of such rapid change is largely unknown. Here, we examined how rarefied species richness, abundance, and composition of amphibians and reptiles varied across forest remnants, cashew orchards and rice paddies in northern Guinea-Bissau. To do so, visual encounter surveys were carried across 21 sampling sites, seven in each habitat type. A total of 703 amphibian and 266 reptile encounters was recorded from nine and 14 taxa, respectively. The results show class-specific responses to habitat type. Amphibian diversity in forest remnants and cashew orchards remained similar, but rice paddies harboured higher abundance and distinct composition compared to forest remnants. Reptile abundance was highest in cashew orchards, which comprised a distinct species composition, when compared to forest remnants. Rice paddies sustained lower reptile richness and abundance. Overall, our results do not support the expected detrimental impacts of cashew expansion, which might be due to the still high heterogeneity of habitat types within the landscape. Rice paddies proved particularly important for amphibians, and for open-habitat reptiles, boosting the landscape-scale species diversity. In face of the eminent habitat conversion, maintaining heterogeneous landscapes, including the persistence of both forest remnants and rice paddies, is critical to minimize biodiversity loss in West Africa.

## 1. INTRODUCTION

Land-use change underpins much of the biodiversity crisis that characterizes the Anthropocene (Dirzo et al., 2014; Newbold et al., 2015; Powers & Jetz, 2019). Today, agriculture occupies ca. 38% of the world’s land surface (FAO, 2020) and as human population grows, this figure is expected to increase (Godfray et al., 2010; Foley et al., 2011). Agricultural-induced land-use change is particularly acute throughout the tropics (Hansen et al., 2020), thus impacting some of the planet’s most biodiverse regions (Dirzo & Raven, 2003). Yet, despite their biodiversity value, the impacts of anthropogenic activities on these ecosystems are disproportionately understudied compared to the temperate regions (Newbold et al., 2020) and most of the research on land-use change in the tropics has been focused on the Neotropics, leaving tropical Africa poorly understood (Gardner et al., 2009)

The impacts of land-use change on biodiversity are diverse, and often taxon-specific (Mendenhall et al., 2014). Yet, the conversion of native habitats to agricultural land typically leads to decreased species richness (Scales & Marsden, 2008), altered species abundance and community composition (Newbold et al., 2015, 2016; Kemp et al., 2019), changed ecological functions (Matuoka et al., 2020) and, ultimately, disrupted ecosystem services (Barnes et al., 2017). The responses of different biological groups to changes in land-use may further vary, and intrinsic species traits make some species more vulnerable than others (Newbold et al., 2014; Rocha et al., 2015). For instance, Harvey & González Villalobos (2007) found birds to be more sensitive to change than bats, as their assemblages varied more across different land-uses, from forests to monocultures. Likewise, Fulgence et al. (2021) observed that Malagasy amphibians exhibited stronger negative responses to land-use change across a gradient from primary forests to agroforests and rice paddies than reptiles.

West Africa is home to a rich biodiversity but has experienced substantial habitat loss and degradation (Lewin et al., 2016), which is anticipated to continue through this century (Powers & Jetz, 2019). Yet, the region has been subject to very few ecological studies and there is a decreasing pattern in the published literature as one moves westward (Luiselli et al., 2019). Guinea-Bissau, on the westernmost tip of the continent and originally covered by a forest-savanna mosaic (Catarino et al., 2008), has much of its native habitats converted to agriculture (Temudo & Abrantes, 2013). Rice (*Oryza glaberrima*) is traditionally cultivated for domestic use, and together with groundnuts comprised the core of the agricultural land in the country until the 20^th^ century (Catarino et al., 2015). After the 1940’s, cashew trees (*Anacardium occidentale*) – native to Northeast Brazil – started to be systematically planted throughout the country, with higher prominence in the north (Temudo & Abrantes, 2014). Cashew orchards – a global agricultural commodity (Rege & Lee, 2023) – have replaced most other forms of land-use in Guinea-Bissau, especially since the 1980’s (Temudo & Abrantes, 2013). Nowadays, agriculture is still the main livelihood in the country, and cashew nuts comprise the only cash crop for the economy of Guinea-Bissau, accounting for 90% of all exports (FAO, 2021).

The once highly complex bio-cultural landscapes in Guinea-Bissau, comprising a forest-rice mosaic known to withstand high biodiversity levels (Temudo et al., 2015), are now threatened by the recent and still ongoing expansion of cashew orchards (Catarino et al., 2015). These are typically dominated by smallholders and are quickly expanding throughout the tropics, homogenizing the landscapes (Rege & Lee, 2023). Notably, these monocultures exhibit a less complex vegetation structure compared to forests (Rege & Lee, 2023; but see Sousa et al., 2015). The true dimension of the impacts of cashew expansion on local biodiversity is little known (Catarino et al., 2015; Monteiro et al., 2017). Yet, the limited available literature regarding species responses to cashew expansion shows declines in species richness and compositional changes across different taxa, compared to the habitats they have replaced (Rege & Lee, 2023). For instance, Komanduri et al. (2023) documented significant shifts in amphibian composition between cashew orchards and forests in India and, in Guinea-Bissau, Vasconcelos et al. (2015) found that butterfly assemblages persisting in cashew orchards were mostly composed of generalist species.

Amphibians and reptiles are among the most threatened animals on Earth (Cox et al., 2022), yet their responses to anthropogenic pressure are less studied than that of other taxa (e.g., invertebrates and birds; Newbold et al., 2014) and there is a strong geographical bias in the available literature, with efforts skewed toward temperate regions and the Neotropics (Guedes et al., 2023; Tan et al., 2023). Although both amphibians and reptiles are ectothermic and thus particularly vulnerable to environmental changes (Newbold et al., 2014; Cordier et al., 2021), the highly permeable skin of amphibians, together with their biphasic life cycle make them particularly sensitive to land-use changes (Winter et al., 2016; Fulgence et al., 2021). To appraise the effects of such land-use changes on these vertebrates, we examined patterns of amphibian and reptile species diversity in forest-cashew-rice landscapes in West Africa. To do so, we assessed species richness, abundance, and composition of both groups within forest remnants, cashew orchards and rice paddies in Northern Guinea-Bissau. Considering the higher structural diversity of forest habitats in contrast to cashew monocultures (Catarino et al., 2015), in addition to the conditions created by the seasonal water availability in rice paddies (Ribeiro et al., 2019), we anticipated that: (1) amphibian and reptile species richness is higher in forest remnants and lower in cashew orchards; (2) abundance of amphibians is higher in rice paddies but that of reptiles is lower; and, consequently, (3) overall species composition is distinct across the different habitat types.

## 2. METHODS

### 2.1 Study area

This study took place in northern Guinea-Bissau, Oio province, around the village of Djalicunda, 8 km south of the city of Farim (12°19’49.82"N, 15°10’57.55"W; Figure 1). In the region, the once forest-savanna mosaic has given way to agricultural land (Catarino et al., 2008), currently consisting of scattered small *tabancas* (villages) surrounded by forest remnants and large areas of extensive smallholder agriculture. These make up mosaics of mostly forest remnants, cashew orchards, and rice paddies. Within this region, cashew orchards are expanding, driving the clearing of some of the last forest remnants (Temudo & Abrantes, 2014). The area has a very smooth relief below 50 m altitude and has defined wet – from June to October – and dry – from October to June – seasons (Catarino et al., 2008). The mean temperature throughout the country ranges between 25.9 and 27.1 °C, and the annual precipitation between 1200 mm in the northeast and 2600 mm in the southwest (Catarino et al., 2008).

**Figure 1.**
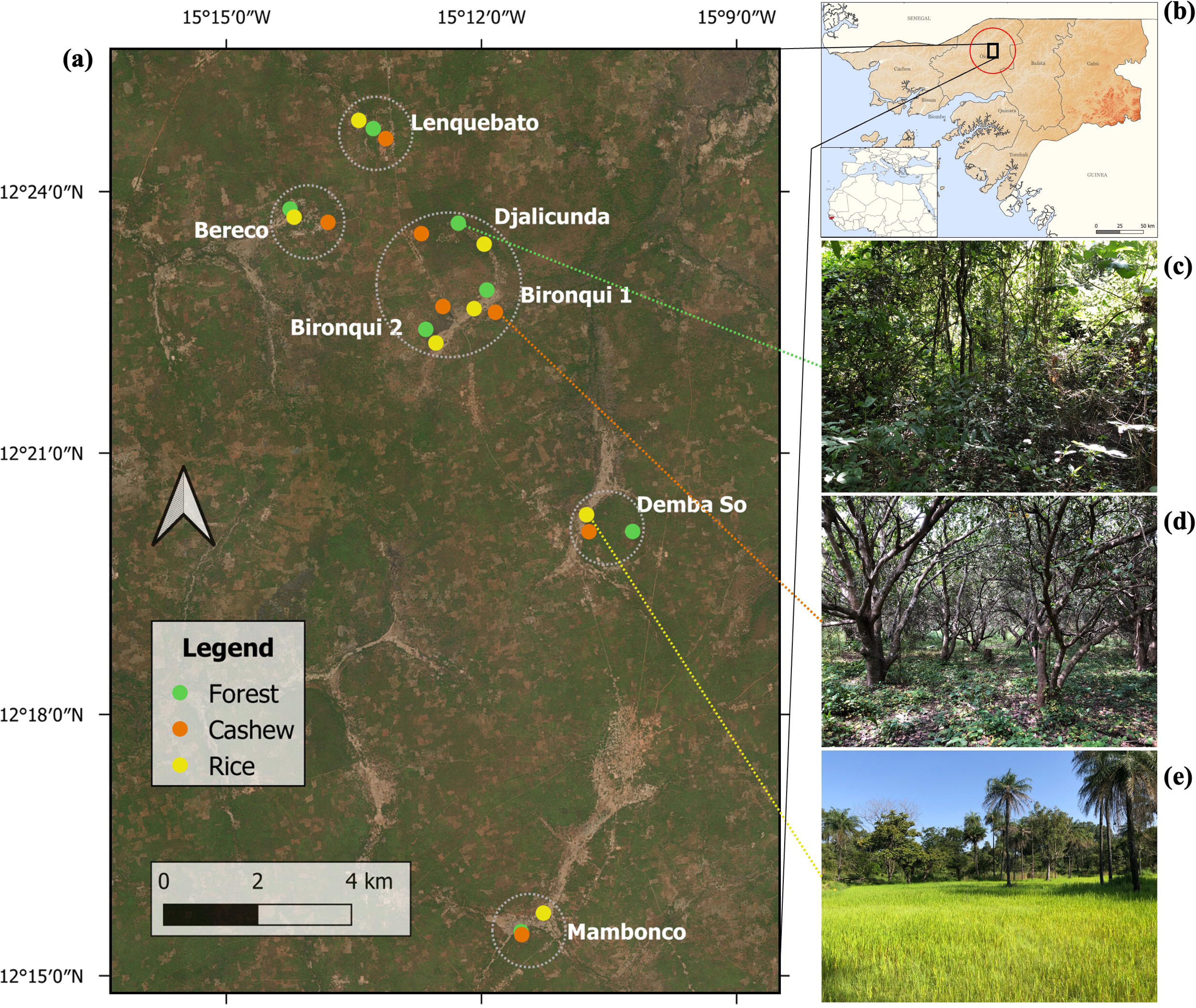
(a) Study area in northern Guinea-Bissau highlighting the location of each of the 21 study sites (solid dots color-coded by habitat type), across seven villages in northern Guinea-Bissau. Dashed grey circles represent the geographically nested structure of the sampling sites; (b) Locations of Guinea-Bissau and of the study area. Each of the sampled habitat types is further illustrated: (c) forest remnants, (d) cashew orchards and, (e) rice paddies (e). Map sources: qGIS (2023), GADM (2021) and geoBoundaries (2017). Photos: Francisco Reis-Silva.

We surveyed amphibians and reptiles across three habitat types: namely, forest remnants, cashew orchards and rice paddies. The surveys took place across 21 study sites, seven of each habitat type and further nested into five locations (Figure 1a; Figure S1). Study sites were selected using rice paddies as a limiting factor, followed by the availability of forest fragments and cashew orchards in the surroundings.

### 2.2 Surveyed habitat types

To describe each study site and understand variation within, but especially between habitat types, we carried a visual characterization from the centre of each site and estimated different metrics, after the rainy season. These included percentage of bare ground, leaf litter, percentage canopy cover, and the number of trees (DBH ≥ 10 cm) and stems (DBH < 10 cm) for a 5 m radius, among others (Table S1). We then used the number of trees to extrapolate the density of trees and stems per hectare for each study site.

*Forest remnants* are mostly second growth open forest subject to variable levels of human intervention (Catarino et al., 2008) for the extraction of forest products, but with little to any tree removal. The canopy cover of forest remnant sites varied between 65 and 95%, and the density of trees between 892 and 1911 trees/ha. Leaf litter ranged between 0 and 90%, and the density of stems varied between 3565 and 16679 stems/ha. Unlike cashew orchards and rice paddies, forest remnants had many thin stems and lianas that increased the overall vertical complexity of the habitat.

*Cashew orchards* had similar arboreal structure to forests yet were of lower height and characterized by the virtual absence of other tree species. There was little to no bare ground (between 0 and 20%). Instead, there were short plants and leaf litter covering the ground. The density of cashew trees within surveyed sites varied between 1656 and 3949 trees/ha, and the canopy was relatively dense (≥80% in six out of the seven sites). Cashew orchards were often crossed by narrow walking paths. The undergrowth is cut short just before flowering, and cashew nut harvesting occurs between June and July. These orchards are biological, as no agro-chemicals nor irrigation are used in their management (Catarino et al., 2015). All cashew orchard sites appeared to have replaced pre-existing forests. The exact age of the orchards is unknown, but since they already produce fruits, they are ≥ 8years old (Dendena & Corsi, 2014).

*Rice paddies* were open areas occupying former seasonally flooded savannas. They were crossed by banks that served as dams and made-up walking paths during the flooded period. There were scattered trees or small groups of trees that may form small islands, with tree density ranging between 0 and 255 trees/ha and almost any canopy cover (between 0 and 5%). These sites typically retain water from the wet season until December. When flooded, the vegetation was made up mainly of rice (*O. glaberrima*), with occasional other plants on the banks and islands. Once rice paddies dry, the ground is covered by low, sparse herbaceous vegetation, with patches of bare ground. Rice is planted in June/July and harvested in November/December, except when the paddy is left fallow. Most of the water available in study area was concentrated in rice paddies.

### 2.3 Herpetofauna surveys

Data collection took place over two four-week long campaigns in 2022. To account for seasonal differences in herptile activity, the first campaign was carried out at the end of the dry season/beginning of the wet season (June/July) and the second one at the end of the wet season (October/November). In each campaign, we surveyed all study sites three times during the day (mostly between 09h15 and 12h30), and once at night (mostly between 19h30 and 22h00), equalling eight surveys at each of the 21 sites.

Herpetofauna surveys took place across 21 circular study sites of 25 m radius in time-standardized surveys (Fulgence et al., 2021). Surveys were carried out by one observer who systematically surveyed the study site for 45 minutes, amounting to a total of 126 sampling hours: 94.5 h during daytime and 31.5 h at night-time. In each survey, the observer thoroughly searched the sites in a zig-zag manner, and carefully checked for herptiles underneath any loose object (e.g., dead wood, bark, leaf litter). We took note of the date, time, and weather at the beginning of every survey. For each amphibian and reptile encounter, we registered the species (or the lowest possible taxonomic level), microhabitat (e.g., tree trunk, leaf litter, under log) and age (i.e., adult or juvenile). At times, photos were used for ID confirmation.

We identified amphibians with the aid of AmphibiaWeb (AmphibiaWeb, 2022) and complementary literature (Pickersgill, 2007; Auliya et al., 2012). For reptile identification, we used Reptile Database (Uetz et al., 2023), and the field guides Chippaux & Jackson (2019) for snakes and Trape et al. (2012) for lizards, crocodiles, and testudines. We identified each herptile down to the lowest possible taxonomic level based on morphological characters. For the 28 times (<3%) we could not identify the specimen to the genus level, we disregarded the encounter (except one record from the Leptotyphlopidae family). Amphibians were analysed at the *taxa* level since five taxa were identified to species and four to genus level, which may include more than one species. Reptiles were assessed at species level, except for the family Leptotyphlopidae represented by one encounter. To streamline, *taxa diversity* is hereafter referred to as *species diversity*.

This work was carried out in cooperation with KAFO, a Guinean NGO that works in close contact with local communities. This organization established the connection between the research team and local communities. The committee of each village was consulted in each field season and granted permission for said work. Herpetofauna surveys were carried out following the appropriate guidelines (Baupre et al., 2004).

### 2.4 Data analysis

For each study site, we summed the number of encounters on all the eight surveys conducted. To assess sampling sufficiency, we made encounter-based species accumulation curves for each of the study sites, using the *rarecurve* function of “vegan” R package (Oksasen et al., 2020). Given limitations in achieving sampling sufficiency for some of the study sites (Figure S1), we used Anne Chao’s proposed method to estimate a rarefied species richness – *Chao1* (Chao, 1987). This metric is often used in the assessment of richness in herpetological studies (e.g., Hutchens & DePerno, 2009; Fulgence et al., 2021) and was obtained using the function *ChaoRichness* from the R package “iNEXT” (Chao et al., 2014; Hsieh & Chao, 2022). We then used three different metrics – rarefied species richness, species abundance and species composition – to examine the effect of habitat type on amphibians and reptiles. Because the functions used for rarefied species richness and composition analyses cannot handle zeros, we removed sites that had no encounters from the subsequent analyses: two sites (Dem-R and Mom-R) from the amphibian, and two (Dem-F and Mom-F) from the reptile analyses. Number of encounters was used as a proxy of species abundance, as been commonly used in other studies (e.g., Fulgence et al., 2021, Komanduri et al., 2023). Species composition was analysed using a Non-Metric Multidimensional Scaling (NMDS) with Bray-Curtis abundance-based dissimilarities through the *metaMDS* function of the R-package “vegan” (Oksasen et al., 2020) (stress = 0.114 and 0.059 for amphibians and reptiles, respectively). Sites Bir1-R, Len-C and Len-R were characterized by only an exclusive species for either of the classes (Len-C had an exclusive amphibian species, and Bir1-R and Len-R exclusive reptile species). Including these discrepant observations in the analysis was leading to NMDS scores different by four orders of magnitude (i.e., Len-C = 5320.9, <0.0; Bir1-R = 2867.5, –996.4; and Len-R = 2353.3, 1189.0). These sites are not shown in the ordination diagram and were removed from subsequent analyses. The scores for the first and second axes of the NMDS were extracted and used as response variables regarding species composition.

We then used Generalized Linear Mixed Models (GLMMs) or Linear Mixed Models (LMMs) to evaluate the effects of the different habitat types on rarefied species richness, species abundance and species composition (NMDS1 and NMDS2). Sites were nested within five landscapes (Figure 1a), so we included the landscape as a random factor to account for the natural variability and distance effects. We checked the distribution of the response variables and fitted appropriate models to the corresponding distributions. As such, we fitted a Poisson distribution (*log* link) for rarefied species richness, and a negative binomial for abundance, as the species abundance residuals were over-dispersed when a model with Poisson distribution (*log* link) was tested. We fitted LMMs with Gaussian distribution for the first and second NMDS axes. Models were computed using the “lme4” package (Bates et al., 2015). All analyses were conducted on the R version 2023.03.0+386 (R Core Team, 2023), and the “ggplot2” R package (Wickham, 2016) was used for visualization.

## 3. RESULTS

We recorded a total of 703 amphibian (77.0% of which were juveniles) and 266 reptile encounters across the 21 sampling sites. Overall, rarefaction curves reveal that amphibians were better sampled than reptiles in the study area (Figure S1), and belonged to nine taxa (five species and four genera), from nine genera and six families; the reptiles to 14 species, from 13 genera and nine families. The most recorded amphibians were *Ptychadena* spp. (54.5%)*, Hyperolius spatzi* (25.0%) and *Leptopelis viridis* (13.8%), while three taxa were only recorded once (0.14%). A total of 622 amphibians were encountered in the rice paddies (88.5%), and all but two amphibian taxa (85.7%) were present in this habitat. Rice paddies also had the greatest number of exclusive amphibian taxa (*Afrixalus vittiger, Hoplobatrachus occipitalis* and *Hildebrandtia ornata*), while four taxa were recorded across all habitat types (*Phrynobatrachus* spp*., L. viridis, Ptychadena* spp. and *H. spatzi*; Figure 2a). Forest remnants and cashew orchards had only one exclusive taxon each (*Hemisus* sp. and *Kassina* sp., respectively), both singletons. The lizards *Trachylepis affinis* (39.1%), *Lygodactylus gutturalis* (37.6%) and *Agama agama* (15.4%) made up most of the reptile records, whereas eight species were recorded only once (0.38%) or twice (0.75%). A total of 179 reptiles were encountered in the cashew orchards (67.3%), and nine out of the 14 reptile species observed were found in this habitat (64.3%). Two reptile species were recorded exclusively in forest remnants (the gecko *Hemidactylus angulatus* and the snake *Atractaspis aterrima*), three in cashew orchards (the gecko *L. gutturalis*, and the snakes *Crotaphopeltis hotamboeia* and *Elapsoidea semiannulata*), and three in rice paddies (the lizards *Latastia ornata* and *Trachylepis perrotetii*, and the cobra *Naja nigricollis*). Only two reptile species were found across the three land-uses (*A. agama* and *Varanus niloticus*; Figure 2b).

**Figure 2.**
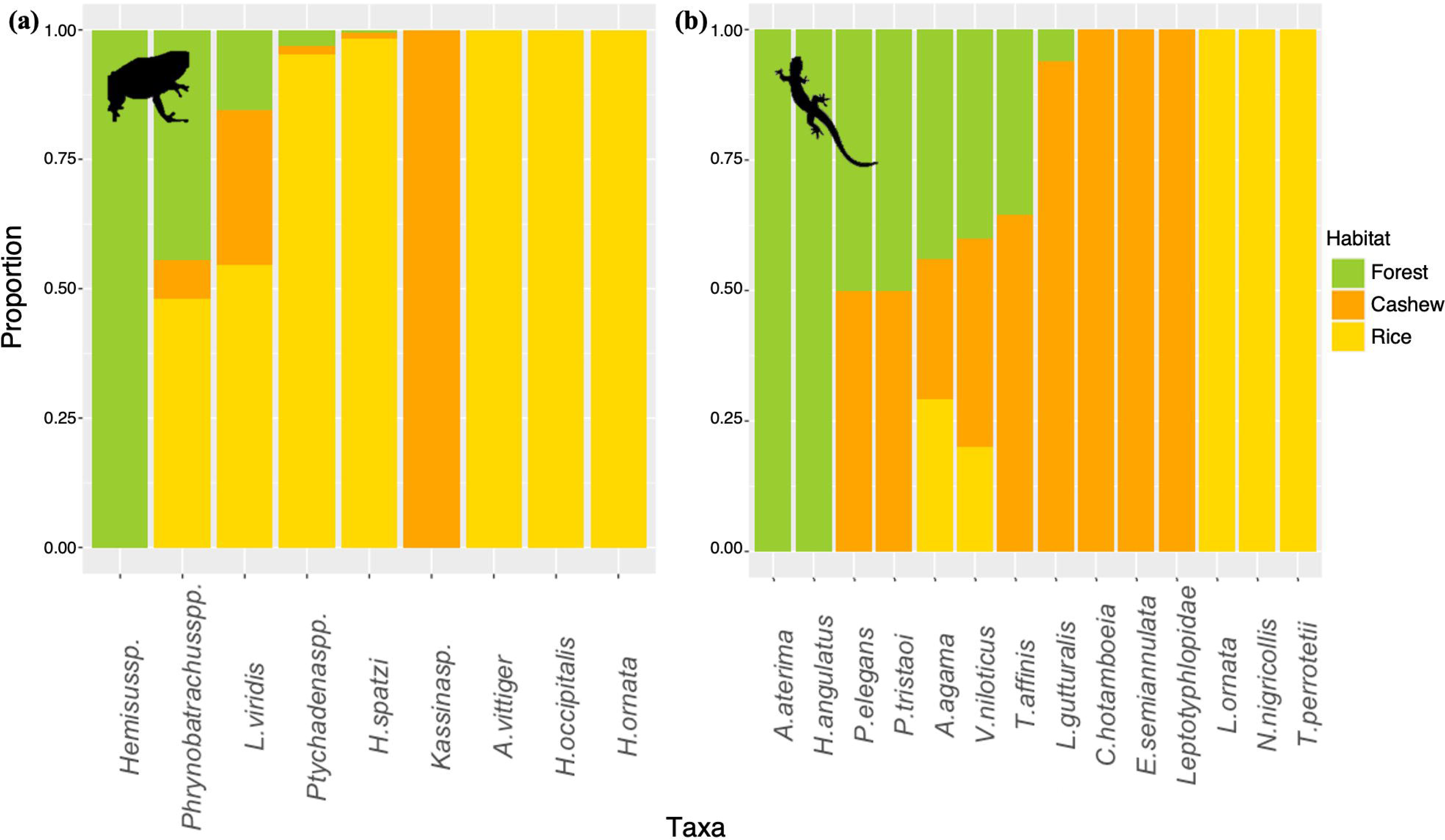
Proportions of (a) amphibian and (b) reptile species encounters across the three sampled habitat types in northern Guinea-Bissau. Data includes nine amphibian (703 encounters) and 14 reptile species (266 encounters).

Amphibian rarefied species richness was similar across habitat types (Figure 3a), but species richness of reptiles was lowest in rice paddies (*β*_habitat_ = –1.301, *P* = 0.002; Figure 3b, Table S2). Amphibian abundance was higher in rice paddies (*β*_habitat_ = 2.843, *P* < 0.0001; Figure 3c, Table S2) than in forest remnants, whereas that of reptiles was higher in cashew orchards (*β*_habitat_ = 0.939, *P* < 0.0001; Figure 3d, Table S2) compared to forest remnants. Yet, it was lower in rice paddies (*β*_habitat_ = –1.415, *P* < 0.0001; Figure 3d, Table S2).

**Figure 3.**
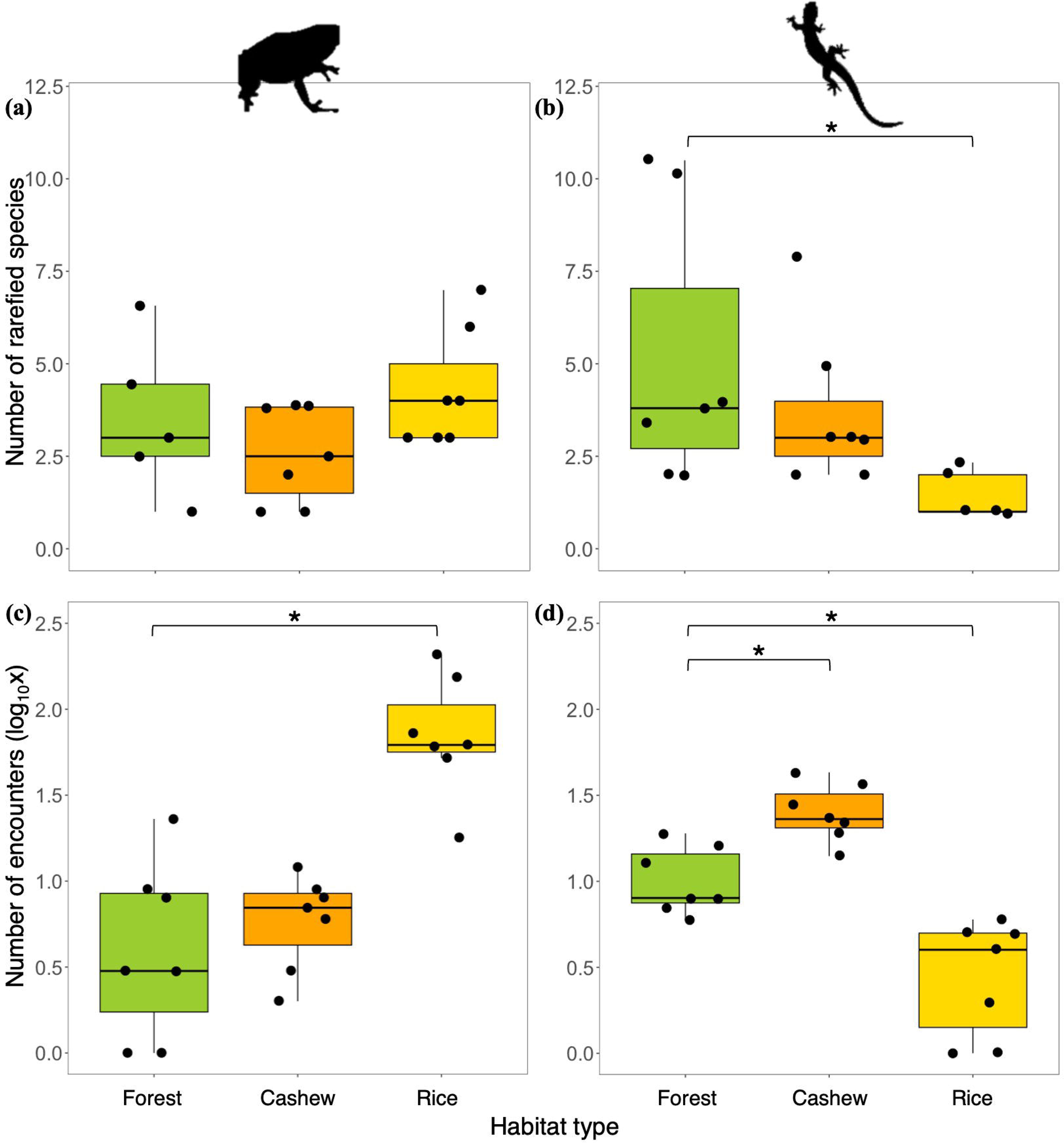
Rarefied species richness (a, b) and observed abundance (log_10_) (c, d) for amphibians (left column) and reptiles (right column), across forest remnants (green), cashew orchards (orange) and rice paddies (yellow) in northern Guinea-Bissau. Two study sites were discarded from amphibian and reptile rarefied richness (two forest remnants and two rice paddies, respectively) due to the absence of records on those sites for each class.

Species composition of amphibians slightly overlapped between forest remnants and cashew orchards, while rice paddies exhibited distinct species composition (Figure 4a). For reptiles, cashew orchards and rice paddy sites form two distinct groups, with rice paddies exhibiting overlap with forest remnants (Figure 4b). Amphibian composition differed between forest remnants and rice paddies (*β*_habitat_ = 1.531, *P* < 0.0001), as denoted from the first axis of the NMDS (Figure 4c, Table S3), but not between forest remnants and cashew orchards. On the other hand, reptile assemblages in forest remnants differed from those of cashew orchards (NMDS1: *β*_habitat_ = –0.761, *P* < 0.015; NMDS2: *β*_habitat_ = 0.673, *P* < 0.01; Figure 4d, Table S2).

**Figure 4.**
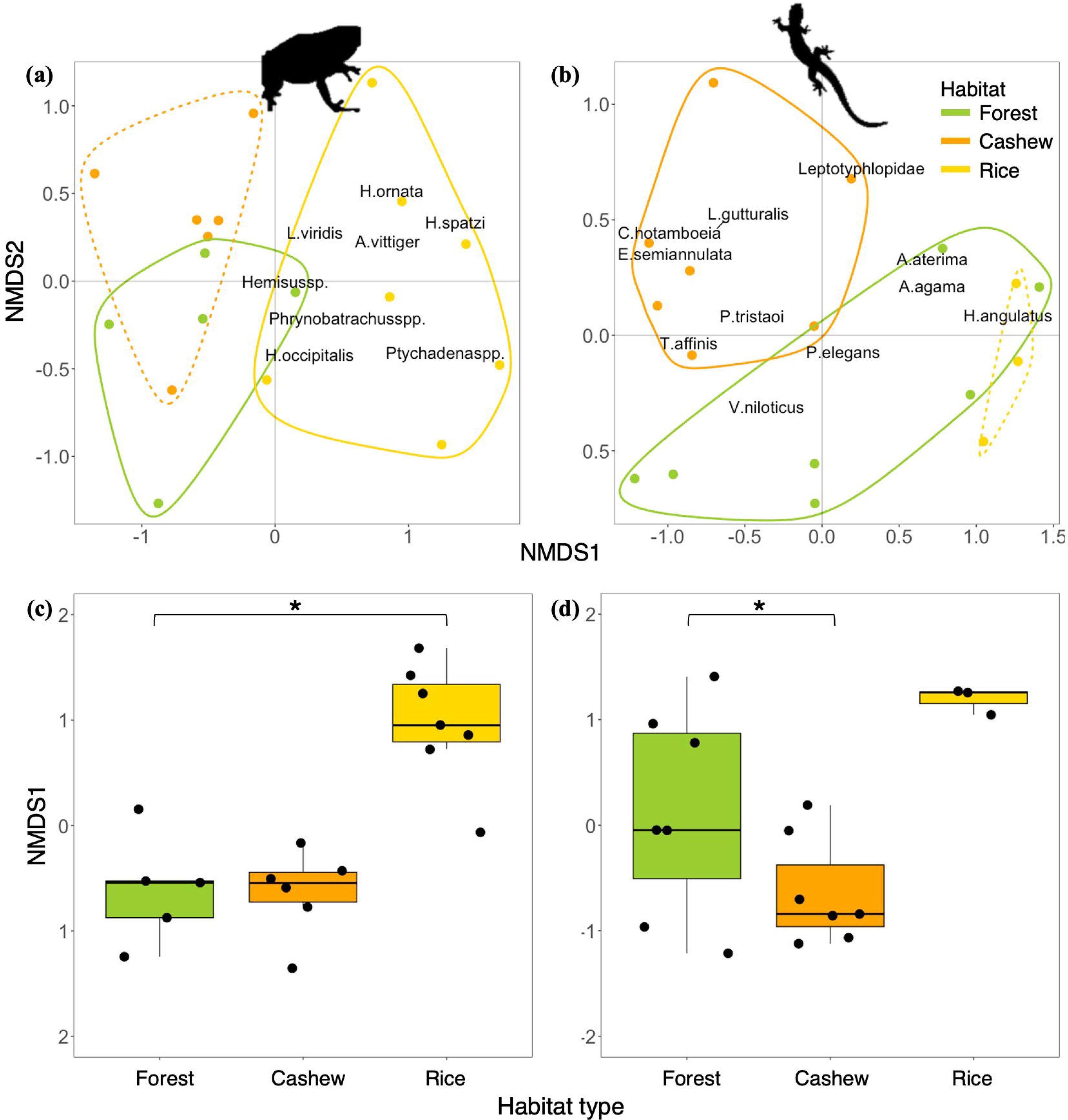
Non-metric multidimensional scaling (NMDS; a, b) of species composition and respective scores for the first axis (NMDS1; b, c) of amphibians (left column) and reptiles (right column) across forest remnants (green), cashew orchards (orange) and rice paddies (yellow) in northern Guinea-Bissau. Points denote study sites and text species; the stress values are (a, c) 0.114 and (b, d) 0.059. Two study sites were discarded from amphibian and reptile composition plots (two forest remnants and two rice paddies, respectively) due to the absence of records on those sites for each class; three other sites were further discarded, as they were outliers – study sites which only species were exclusive to them (a, c) one cashew orchard and (b, d) two rice paddies); the dashed lines

## 4. DISCUSSION

In the context of the rapid ongoing land-use changes in West Africa, here we provide the first insights on how amphibians and reptiles can cope with such changes. Overall, we found that both groups respond differently to land-use, with amphibians being particularly diverse in the rice paddies, whereas reptiles presented higher abundance in the cashew orchards. Forest habitats did not stand as sites of higher herpetofauna diversity, making our findings not agreeing with initial expectations in that regard.

### 4.1 Responses to cashew expansion

All the diversity metrics of amphibians and the rarefied species richness of reptiles remained similar across habitat types. Although this was an unexpected result, it might be linked with (1) the structural similarity between cashew orchards and native forest, higher than initially anticipated especially given the low habitat quality the remaining forests and with (2) the high habitat heterogeneity characterising the study area (Estrada-Carmona et al., 2022). Human-modified habitats that retain the vertical structure of pre-existing vegetation are typically important for maintaining species diversity in agricultural landscapes (Newbold et al., 2014). Both forest remnants and cashew plantations in our study area are characterized by having substantial ground cover, a high density of trees and a dense canopy cover that filters sunlight and helps with temperature regulation within (Table S2). In addition, cashew orchards are poorly intensively manged and not subject to substantial application of chemical products. This possibly justifyies the similarities in herptile assemblages between cashew and forest remnants (but see Vasconcelos et al., 2015). Similarly, in the Western Ghats (India), Komanduri et al. (2023) found that cashew orchards to have similar amphibian abundances to those in forests, and to withstand 67% of the amphibian species that occur in the region. Also in India, a subset of terrestrial mammals in the forest also makes use of cashew plantations (Rege et al., 2020), while bird diversity in cashew plantations was comparable to that of adjoining forests (Munge & Kumar, 2022). Moreover, in the case of northern Guinea-Bissau, remaining forests are of relatively small size and subject to regular human intervention, as those correspond to community-managed forests (e.g., non-timber forest products extraction, Palmeirim et al., 2023), and thus may no longer withstand the species they once did, as habitat specialists may have gone extinct (Devictor et al., 2008; Palmeirim et al., 2017). Yet, due to lack of reference data, it is impossible to know the species these forests supported. Furthermore, our study area is characterized by a high degree of habitat heterogeneity, which is also clear by the relatively short distances between sites of different habitat types (see Figure 1). Usually, given the higher diversity in habitat types per unit of area, heterogenous landscapes tend to harbour higher biodiversity levels than homogenous ones (Bhagwat et al., 2008; Estrada-Carmona et al., 2022). In sum, although our results seem to suggest that cashew orchards can withhold high herptile diversity compared to forest, this might reflect impoverished forest assemblages, not representing a suitable baseline. Indeed, we would expect to have been able to record several additional species (e.g., the toad *Sclerophrys ssp.*, the chameleon *Chamaeleo gracillis;* Chippaux & Jackson, 2019; Trape *et al*., 2012; AmphibiaWeb, IUCN, 2022; Uetz *et al*., 2023). One caveat to this, however, was that our sampling effort that was not always enough to obtain representative species assemblages for all studies sites (see Figure S2).

Notwithstanding the similar patterns of overall species diversity between forests and cashew orchards, the latter tended to be dominated by habitat generalists, that are widely distributed throughout Sub-Saharan Africa. These included the frogs *L. viridis*, *Ptychadena* spp. and *Phrynobatrachus* spp. (Nowakowski et al., 2017; AmphibiaWeb, 2022; IUCN 2022), which made up 92.5% of amphibian encounters in cashew orchards, and the reptiles *T. affinis* and *L. gutturalis*, which encompass 90.0% of the observations in this habitat type (Trape et al., 2012; IUCN, 2022). Likewise, in a study also carried out in the North of Guinea-Bissau, Vasconcelos et al. (2015) found that cashew orchards were mostly occupied by generalist butterfly species. It is possible that similarities in lizard richness between habitat types are in part due to specialists being replaced by generalist species (Palmeirim et al., 2017).

Moreover, the higher species abundances of reptiles in cashew orchards were due to the increased number of encounters of the generalist species, namely *T. affinis* and *L. gutturalis*. Herptile species persisting in human-modified landscapes often exhibit higher abundances in the altered than in native habitats, as explained by the success of those species in making use of the trophic, structural, and climatic resources available (Newbold et al., 2014). Nevertheless, the lower degree of obstruction in cashew orchards might have favored species detectability. This limitation urges caution when interpreting the results on this regard.

### 4.2 The role of rice paddies

Rice paddies hold conspicuously greater amphibian abundance than the remaining habitat types, while the opposite trend was noted for reptiles. The higher abundance of amphibians in rice paddies likely reflects the breeding ecology of these animals, which depend on water for reproduction (Semlitsch et al., 2015; Ribeiro et al., 2019; Fulgence et al., 2021). The particularly high abundance is further due to the typical *r*-strategist of amphibians, as noted given that 77.0% of the recorded amphibians were juveniles. On the other hand, the reduced diversity of reptiles found in rice paddies aligns with similar findings from Madagascar (Fulgence et al., 2021). Before the rainy season, rice paddies are dry and with little cover on the floor. After the start of the rain, rice paddies become flooded which, despite beneficial for amphibians, makes this habitat less suitable for terrestrial reptiles. The little non-flooded area, including shade and shelter availability, make rice paddies only able to sustain species-poor reptile assemblages (Fulgence et al., 2021).

Amphibian species composition in rice paddies differed substantially from forest and cashew orchards. This seems to reflect both the presence of exclusive taxa, and the high amphibian abundance in that habitat. Our results seem to indicate that reptile composition in rice paddies represents a subgroup of that of forest remnants. However, this might reflect the exclusion of four rice paddy sites (out of seven) from the composition analysis: only *Agama agama* and *Varanus niloticus* were included for the rice paddies (both also recorded in forest remnants). Yet, each class had three exclusive taxa in rice paddies: the frogs *Hoplobatrachus occipitalis, Afrixalus vittiger* and *Hildebrandtia ornate*; and the cobra *N. nigricollis*, the skink *T. perrotetii* and the lizard *L. ornata* (Fig. S2). Except for *N. nigricollis*, a habitat generalist, all these are savanna-dwelling species (Trape *et al*., 2012; Chippaux & Jackson, 2019; AmphibiaWeb, 2022). Noteworthy, this was the first time *L. ornata* was observed in Guinea-Bissau since the collection of the type specimen in 1938, and represents the third ever report of the species (Monard, 1940; Pauwels et al., 2023). The presence of these species, together with the high amphibian abundance suggests that rice paddies have an important conservation value for the herpetofauna of in Guinea-Bissau. Indeed, in addition to complementing the different habitat requirements amphibians have throughout their life cycle (Ribeiro et al., 2019), the rice paddies seem to be somewhat structurally analogous to the savannas they have replaced and retain at least some of the open-habitat species of the forest-savanna mosaic.

### 4.3 Conservation implications

The different responses of amphibians and reptiles to cashew orchards and rice paddies emphasizes the importance of assessing each separately (Cordier et al., 2021). Yet, despite following a similar study design to that of Fulgence et al. (2021), our work does not support the premise that amphibians are more sensitive to land-use change than reptiles. In fact, while amphibian diversity only varied across habitats for two of the assessed metrics (species richness and composition), reptiles showed variation in all three assessed diversity metrics.

Rice paddies are of high importance for maintaining amphibian diversity in the mosaic landscapes characterizing the northern part of Guinea-Bissau. At the same time, reptiles do better in closed-canopy habitats, forests and cashew orchards. We suggest that the mosaic made by both open-area and closed-canopy habitats is conserved, allowing to maximize the diversity of herpetofauna at landscape level. Alarmingly, particularly forests are quickly being replaced by vast areas of cashew orchards (Temudo & Abrantes, 2014). The fact that forest remnants appear to have comparable diversity to an agricultural habitat must be looked at carefully, and the limitations of this study considered. The importance of rice paddies at landscape level seems to be clearer: in addition of acting as an important breeding place for amphibians, rice paddies seem to act as surrogate habitats to the lost savanna and maintain important herptile assemblages. From an ecosystem-service point of view, the presence of amphibians in high abundances in rice paddies may also be a relevant pest-control agent (Hocking & Babbitt, 2014). In face of the imminent habitat conversion, our work suggests that cashew cultivation, and the economical and societal benefits it entails (Dendena & Corsi, 2014), may be possible if maintaining a heterogeneous landscapes. As such, the land occupied by cashew orchards should be balanced against that occupied by rice paddies and native forest remnants.

## Acknowledgements

We thank the members of the GCC Group (University of Helsinki) who gave crucial advice, especially Federica Manca and Tiago Monteiro-Henriques, who helped during statistical analyses; and Amaia Gonzaga Roa and Aina Rossinyol Fernàndez, who helped producing the maps. We are grateful to the workers of the NGO KAFO (Guinea-Bissau), including Ami, Belomi, Djunco, Djari, Francisco, Jara, Judite and Ioba; to the people of Djalicunda and all the *tabancas* where our fieldwork was conducted. We acknowledge financial support from *EcoPestSupression* (cE3c, Lisbon, Portugal; reference no. PTDC/ASP-AGR/0876/2020) and *TROPIBIO* (CIBIO, Vairão, Portugal; European Union’s Horizon 2020 research and innovation programme under grant agreement no. 854248).

## Author contributions

FRS, RR, AR and AFP conceived and designed the methodology. SS settled the logistics required for the field work. FRS collected the data, analyzed the data led the writing. All authors contributed critically to the drafts and gave final approval for publication.

## Notes

### Competing Interest Statement

The authors have declared no competing interest.

