## Supplementary Material for "Unveiling how herpetofauna cope with land-use changes— insights from forest-cashew-rice landscapes in West Africa"

This supporting information includes Table SX and Figures SX to SX

**Table S1.** Habitat characterization of the 21 sampling sites, seven of each habitat type - forest remnants, cashew orchards and rice paddies - after the rainy season in northern Guinea-Bissau.

| Site | bare ground (%) | leaf litter (%) | grass cover (%) | tall grass cover (%) | shrub cover (%) | mid-canopy cover (%) | upper-canopy cover (%) | clutter (0-3) | no. of trees DBH >10cm | Average height 5 highest | termite mounds (0-3) | wood piles/fallen trees (0- | water cover (%) | habitat |
|---|---|---|---|---|---|---|---|---|---|---|---|---|---|---|
| **Ber-C** | 5 | 0 | 85 | 5 | 5 | 85 | 0 | 1 | 16 | 6 | 2 | 1 | 0 | C |
| **Ber-F** | 40 | 10 | 5 | 5 | 40 | 95 | 5 | 2 | 7 | 25 | 1 | 2 | 0 | F |
| **Ber1-R** | 5 | 0 | 60 | 5 | 0 | 5 | 0 | 0 | 0 | 5 | 1 | 3 | 15 | R |
| **Bir1-C** | 5 | 45 | 45 | 5 | 0 | 100 | 0 | 1 | 14 | 6 | 1 | 1 | 0 | C |
| **Bir1-F** | 50 | 0 | 10 | 20 | 20 | 90 | 0 | 1 | 13 | 10 | 1 | 1 | 10 | F |
| **Bir-R** | 0 | 0 | 0 | 70 | 0 | 5 | 5 | 0 | 1 | 1 | 0 | 0 | 65 | R |
| **Bir2-C** | 20 | 20 | 50 | 10 | 0 | 90 | 0 | 1 | 20 | 6 | 0 | 2 | 0 | C |
| **Bir2-F** | 10 | 5 | 40 | 40 | 5 | 70 | 10 | 1 | 12 | 22 | 2 | 2 | 0 | F |
| **Bir2-R** | 5 | 0 | 5 | 60 | 0 | 0 | 0 | 0 | 0 | 9 | 1 | 1 | 45 | R |
| **Daj-C** | 0 | 25 | 65 | 10 | 0 | 90 | 0 | 1 | 13 | 13,6 | 2 | 0 | 0 | C |

| | | | | | | | | | | | | | | |
|---|---|---|---|---|---|---|---|---|---|---|---|---|---|---|
| **Dem-C** | 10 | 5 | 60 | 20 | 5 | 65 | 0 | 1 | 24 | 22 | 1 | 2 | 0 | C |
| **Dem-F** | 0 | 90 | 5 | 5 | 0 | 95 | 5 | 3 | 11 | 20 | 2 | 1 | 0 | F |
| **Dem-R** | 0 | 0 | 0 | 80 | 5 | 0 | 0 | 0 | 0 | 15 | 1 | 0 | 85 | R |
| **Dja-F** | 5 | 5 | 30 | 20 | 40 | 70 | 15 | 2 | 15 | 25 | 1 | 1 | 0 | F |
| **Dja-R** | 5 | 0 | 0 | 30 | 0 | 0 | 0 | 0 | 0 | 8,4 | 1 | 1 | 65 | R |
| **Len-C** | 5 | 20 | 70 | 5 | 0 | 95 | 0 | 0 | 31 | 6 | 2 | 1 | 0 | C |
| **Len-F** | 25 | 20 | 30 | 15 | 10 | 90 | 5 | 2 | 12 | 21 | 1 | 1 | 0 | F |
| **Len-R** | 5 | 0 | 5 | 50 | 0 | 0 | 5 | 0 | 2 | 11,8 | 1 | 0 | 70 | R |
| **Man-C** | 0 | 5 | 50 | 30 | 15 | 90 | 0 | 1 | 15 | 6,4 | 1 | 1 | 0 | C |
| **Man-F** | 5 | 80 | 5 | 5 | 5 | 95 | 5 | 3 | 15 | 18 | 2 | 1 | 0 | F |
| **Man-R** | 0 | 0 | 85 | 5 | 0 | 5 | 0 | 0 | 0 | 4 | 1 | 0 | 90 | R |

8

**Table S2.** Summary of Generalized Linear Mixed Models (GLMMs) and Linear Mixed Models (LMMs) explaining rarefied ('*Chao1*') species richness (GLMMs), observed ($log_{10}x$) abundance (GLMMs) and NMDS axes 1 and 2 (LMMs) for amphibians and reptiles across three habitat types from 21 study sites in northern Guinea-Bissau; t-values are shown for models fitted with a Gaussian distribution (NMDS axes 1 and 2), and z-values for models fitted with a Poisson (rarefied species richness) and negative binomial (species abundance). As an exception, the models for rarefied richness of both classes were based on 19 sites, as two sites for each class had no records. Significant p-values are denoted in bold.

| Response variable | Class | Model parameters | Estimate | Std. error | t/z-value | p-value |
|---|---|---|---|---|---|---|
| **Rarefied species richness** | Amphibians | Forest (Intercept) | 1.253 | 0.278 | 4.501 | **<0.0001** |
| | | Cashew | -0.274 | 0.332 | -0.823 | 0.410 |
| | | Rice | 0.183 | 0.302 | 0.607 | 0.544 |
| | Reptiles | Forest (Intercept) | 1.637 | 0.117 | 9.826 | **<0.001** |
| | | Cashew | -0.325 | 0.257 | -1.264 | 0.206 |
| | | Rice | -1.301 | 0.413 | -3.150 | **0.002** |
| **Species abundance ($log_{10}x$)** | Amphibians | Forest (Intercept) | 1.434 | 0.421 | 3.403 | **<0.01** |
| | | Cashew | 0.084 | 0.445 | 0.188 | 0.851 |
| | | Rice | 2.843 | 0.428 | 4.642 | **<0.0001** |
| | Reptiles | Forest (Intercept) | 2.303 | 0.173 | 13.287 | **<0.0001** |
| | | Cashew | 0.939 | 0.227 | 4.143 | **<0.0001** |
| | | Rice | -1.415 | 0.323 | -4.376 | **<0.0001** |
| **NMDS 1** | Amphibians | Forest (Intercept) | -0.632 | 0.249 | -2.236 | **<0.05** |
| | | Cashew | -0.105 | 0.204 | -0.516 | 0.6060 |
| | | Rice | 1.531 | 0.193 | 7.948 | **<0.0001** |
| | Reptiles | Forest (Intercept) | -0.001 | 0.310 | -0.031 | 0.976 |
| | | Cashew | -0.761 | 0.311 | -2.445 | **0.015** |
| | | Rice | 0.800 | 0.417 | 1.921 | 0.055 |
| | Amphibians | Forest (Intercept) | -0.327 | 0.272 | -1.201 | 0.230 |

| | | | | | | |
|---|---|---|---|---|---|---|
| **NMDS 2** | | Cashew | 0.643 | 0.368 | -1.747 | 0.081 |
| | | Rice | 0.289 | 0.356 | 0.811 | 0.416 |
| | | Forest (Intercept) | -3118 | 0.156 | -1.994 | **<0.05** |
| | Reptiles | Cashew | 0.673 | 0.221 | 3.044 | **<0.01** |
| | | Rice | 0.196 | 0.285 | 0.687 | 0.492 |

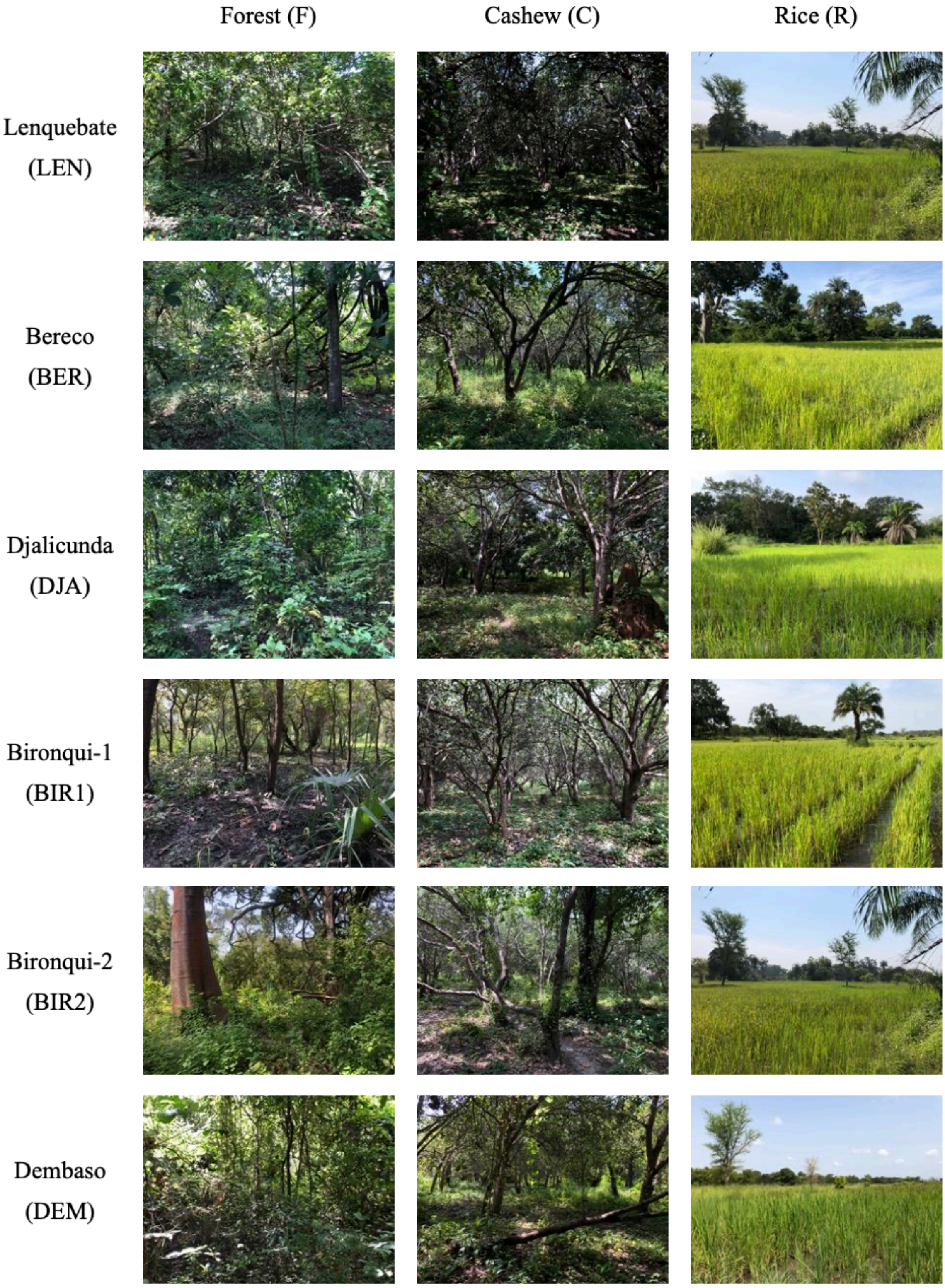


**Figure S1.** Photo of each of the 21 sampling sites, including three sites of each habitat type: forest remnants, cashew orchards and rice paddies across seven *tabancas* (villages) in northern Guinea-Bissau. Photo credits: Francisco dos Reis-Silva. Lines indicate sites within the same village and columns indicate the different habitat types. Abbreviations as used in the main text are further indicated.

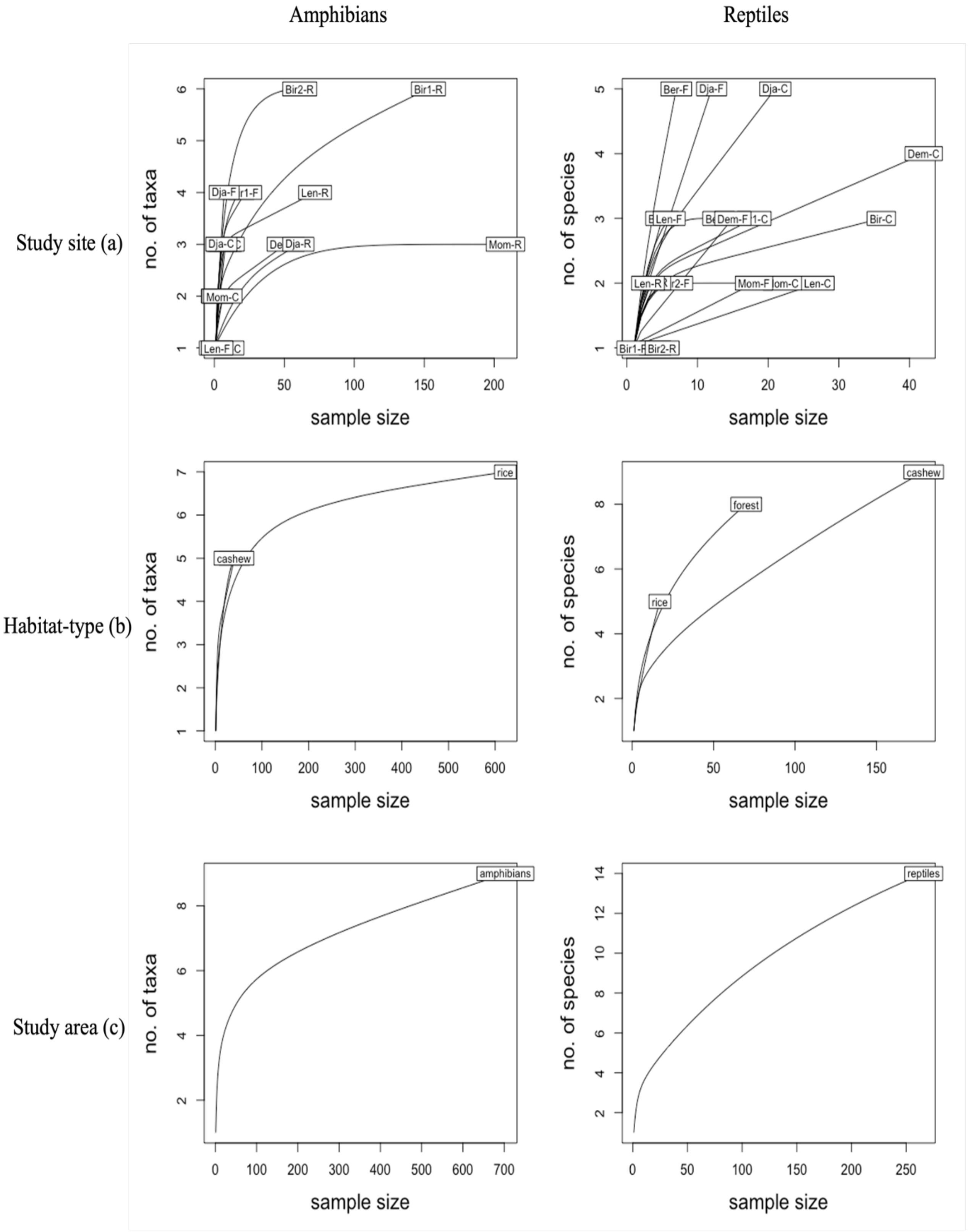


**Figure S2.** Encounter-based species accumulation curves for amphibians and reptiles in north Guinea-Bissau across (a) sampling sites, (b) habitat type and (c) study area.

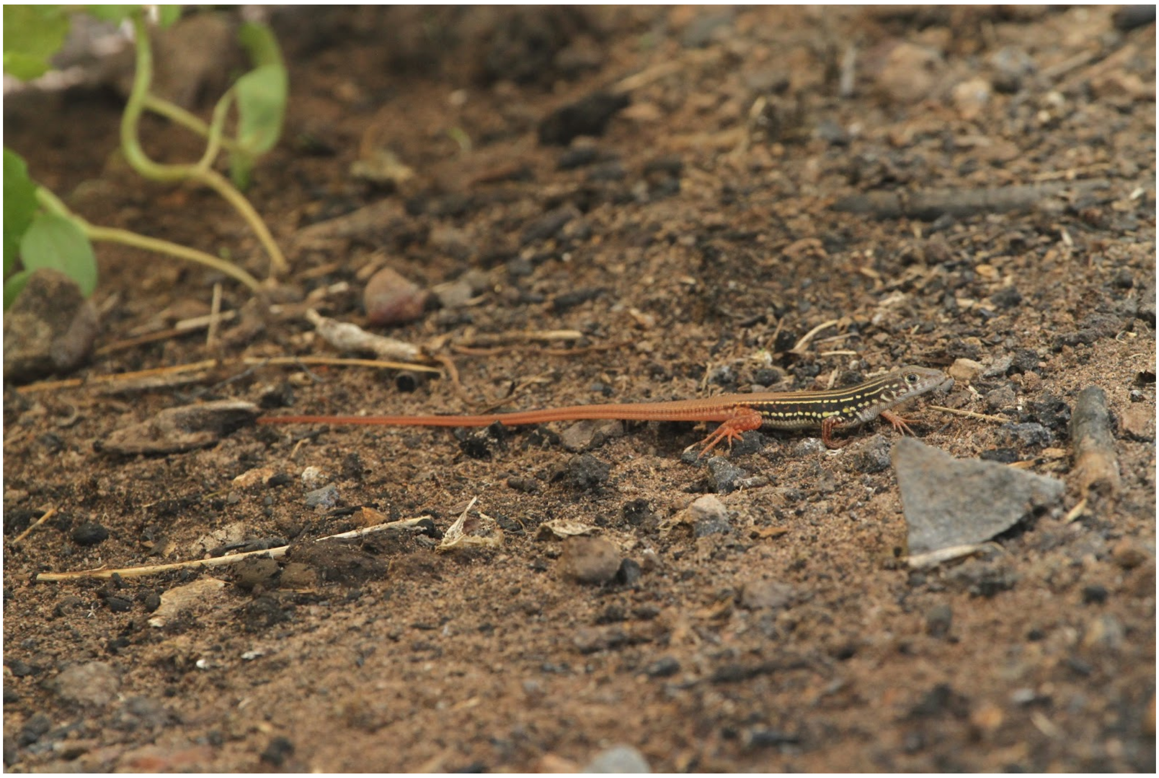

**Figure S3.** *Latastia ornata in situ.* First record of *Latastia ornata* since the collection of the type specimen in 1938, and third ever documented record of the species. Photo credits: Ricardo Rocha
